# Rats pursue food and leisure following the same rational principles

**DOI:** 10.1101/2024.12.08.627420

**Authors:** Raegan S. Logue, Ana K. Garcia, Brissa A. Bejarano, Taylor Fujioka, Mikayla Duke, Riley K. Kendall, Andrew M. Wikenheiser

## Abstract

Animals in the wild must balance multiple, potentially mutually-exclusive goals simultaneously in order to survive. Yet laboratory tests of decision making often investigate how animals optimize their behavior to achieve a single, well-defined goal, which is often a nutritive reward. Thus, how animals solve multi-objective optimization problems is not well understood. Here, we devised an ethologically-inspired decision making task to examine how rats balance the pursuit of food and non-food reinforcement. Rats performed a free-choice patch-foraging task, in which they could earn food in one location (food patch) or interact with a rodent play structure in a different location (toy patch). The cost of switching between patches was manipulated by requiring rats to endure a long or short “travel time” penalty during which they were not able to access either patch. Rats devoted a considerable amount of their limited foraging time to patches of both types, showing a small but significant preference for food patches. In accordance with theoretical models of foraging, when the cost of switching patches was high rats chose longer stay durations in both types of patches, suggesting that similar rational principles guided their pursuit of food and non-food rewards. Examining the within-session dynamics of time allocation revealed that rats showed an early preference for spending time in toy patches that reversed over the course of the session. Satiety manipulations demonstrated that patch residence time decisions were under goal-directed control, and responsive to current needs and recent consumption. These results validate a naturalistic approach to testing decision making in rats over a range of food and non-food goods. Key words: Patch foraging, decision making, multi-objective optimization, play, leisure

**Significance Statement:** The mechanisms by which animals trade off competing goals remain poorly understood. Here, we investigated whether rats use rational decision-making strategies to balance essential needs with non-essential pursuits like leisure by letting rats choose between earning food and engaging with a play structure. Rats performing a free-choice patch-foraging task adjusted their behavior based on opportunity costs, staying longer in patches when switching costs were high, in accordance with models of optimal foraging. Rats allocated significant time to toy patches, even at the expense of food rewards, underscoring the intrinsic value of non-food reinforcers. These results validate a naturalistic approach for studying tradeoffs between qualitatively-different rewards, and advance our understanding of multi-goal optimization in the context of a naturalistic decision problem.

## Introduction

Animals in the wild must manage the pursuit of many goals simultaneously. Goals differ across species, but may include finding food, protecting territory, pursuing mates, and caring for offspring—all while avoiding predators (1, 2). In times of relative abundance, and absent heavy threat of predation, some species also divide their time between work-like and leisure-like activities including rest and play (3, 4). Balancing these competing demands requires organisms to make tradeoffs, prioritizing different goals at different times depending on their current biological needs and the state of the environment (5–9). For most organisms, survival therefore depends not on optimizing behavior to achieve any single goal, but rather adaptively compromising to pursue a range of ends whose costs and benefits vary over time (10). Understanding how animals prioritize competing goals is important for understanding learning and decision making at the behavioral, computational, and neural levels (11–14).

In contrast, laboratory tests of decision making typically ask animals to solve a single, well-defined problem in exchange for a single type of programmed reinforcer, often a nutritive food or liquid. This framework has revealed much about the neural and behavioral mechanisms of decision making, but is not well-suited to investigating how animals balance the pursuit of multiple, competing goals. Nevertheless, there is evidence that even in the context of a single focal decision-making problem animals optimize over multiple distinct objectives beyond the reinforcer programmed by the experimenter. For instance, animals frequently intersperse bouts of steady task performance with breaks, during which they rest, groom, or explore their surroundings. Economic and reinforcement learning frameworks have been used to model breaks in task performance as a simple form of leisure (15–19), which may offer utility by reducing effort costs and attentional demands while also freeing up time for subjects to engage in other potentially rewarding behaviors. However, the typically impoverished set of alternative reinforcers available in a standard operant box make it difficult to ascertain whether breaks in task performance are caused by a positive attraction to some alternative option or processes like boredom or fatigue induced by the focal task.

Foraging theory has proven to be a successful framework for understanding a wide range of ethologically-relevant behaviors, and has shaped how biologists and neuroscientists conceptualize decision making (2, 20). Classical foraging models are based heavily on the principle of lost opportunity: every decision in favor of one course of action carries a cost in terms of alternative sources of reward that cannot also be pursued (21, 22). Although foraging models were originally formulated to explain food-seeking behavior, minimizing opportunity cost is important whenever limited time or effort must be allocated among competing goals. Yet foraging theory has seldom been extended to decisions involving non-nutritive reinforcers and it is therefore unclear whether the same cost-benefit tradeoffs animals are sensitive to when making food-choice decisions also govern behavior in the pursuit of non-food goods.

It is well known that the nature of a reinforcer can influence the form of behavior it motivates. For instance, Petrinovich and Bolles (23) found that food and water deprivation states engaged different behavioral repertoires, with hunger promoting variability and exploration, while thirst elicited more stereotyped responses. Similarly, different cues become preferentially linked with different reinforcers, suggesting that the nervous system processes cue-outcome pairings in a way that depends on their ecological relevance (24). These findings raise the question: do animals apply a unitary foraging strategy across diverse goals, or do different reward types engage distinct strategies?

Here, we tested how rats chose to spend a limited time budget in pursuit of two qualitatively-distinct forms of reinforcement. We offered hungry rats a choice between earning food reward in one location or engaging with a playground-like toy apparatus in a different location. The reward schedule for the food option was fixed, which made the opportunity cost of engaging with the toy structure well defined and easily computed as the amount of food rats would have earned had they eschewed the toy option in favor of pursuing food. By varying the time penalty associated with switching between the food and toy options, we investigated whether rats’ consumption of either or both options responded rationally to changes in the opportunity cost of time.

## Results

### A free-choice foraging task for comparing food and non-food reward

Long-Evans rats (N = 10, 5 female) were tested on a three-chamber apparatus, consisting of two open-field arenas connected to one another by a corridor with computer-controlled doors (**Figure 1A**). Rats were placed in the corridor and the session began when both doors opened simultaneously, allowing them to choose an arena to enter. Rats were free to stay in the arena they selected for as long as they wished. Upon entering one arena, the door to the unchosen arena closed, preventing rats from transiting directly between arenas. To switch between arenas the rat entered the travel corridor, which caused the door to the previously-occupied arena to close, confining the rat to the corridor for a fixed duration to simulate travelling to a new foraging location. After each travel duration, the doors to both arenas opened simultaneously as before. Behavioral sessions continued in this way for 30 minutes. This task comprises a free-choice version of the patch-leaving problem, which has been studied extensively in behavioral ecology, psychology, and neuroscience (21, 22, 25–32).

**Figure 1.**
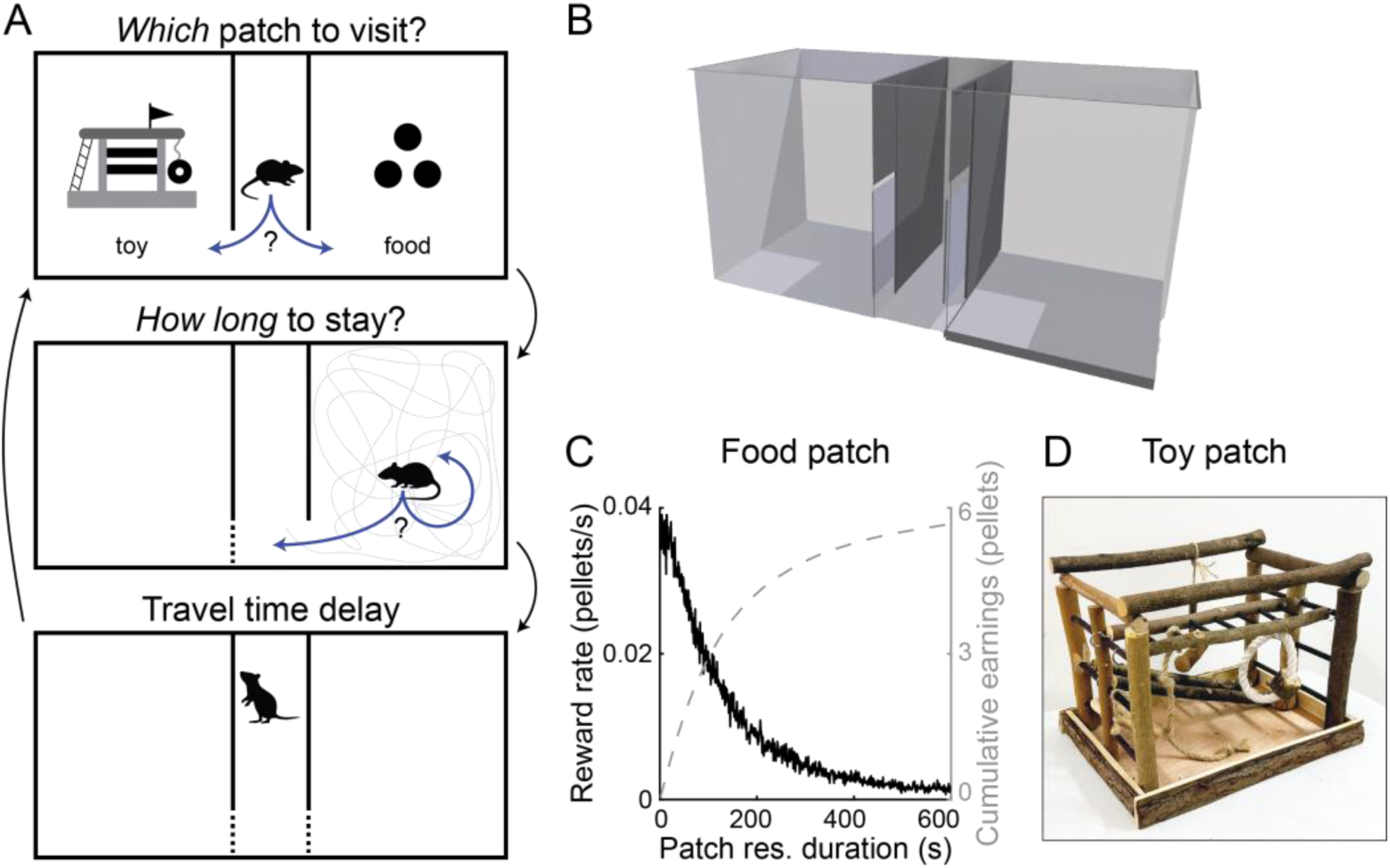
The free-choice foraging task. (A) A cartoon schematic depicts the structure of the free-choice foraging task. Rats began each session in the travel corridor, and first selected which of two foraging patches to visit. After choosing one, they decided how long to remain in that patch. Returning to the travel corridor ended the visit to a patch, and triggered the travel time delay. After the travel time elapsed, rats were again free to select a patch to forage in. This sequence continued for the duration of behavioral sessions. Automated doors ensured that rats could not bypass the travel corridor and pass directly between the foraging patches without experiencing a travel time delay. (B) Three-dimensional rendering of the foraging task apparatus. Central walls (depicted in grey) separated the two patches and defined the travel corridor that rats passed through to switch between locations. Motorized doors (depicted in blue) controlled rats’ access to the patches. (C) When rats visited food patches, sucrose pellets were dispensed into the arena at a rate that decreased over time. The reward rate was maximal when rats first entered the patched, decreased over the course of their visit, and reset to the maximal rate on each subsequent visit to the patch. (D) In toy patches, rats encountered a rodent play structure in the center of the patch, and were free to interact with it.

In each session the arenas were designated as food or toy patches. In food patches, sucrose pellets were dropped from above while rats visited the arena; pellets scattered unpredictably and approximately uniformly across the arena. The reward rate decreased over time during each visit (**Figure 1B**), but reset to the same initial rate when rats re-entered a food patch after a travel delay. In toy patches, a structure consisting of climbing platforms, tunnels, and fixed, manipulable objects (strings, wood blocks, etc.) was positioned in the center of the arena (**Figure 1C**) and rats were free to interact with it. Food was not available in toy patches. Rats experienced a randomized sequence of eight session types, which fully crossed patch type (food or toy) and arena location (left or right) at two levels of travel time (60 s or 10 s). Rats were food restricted for the duration of the experiment.

### Rats balanced pursuit of food and non-food reward

Rats visited toy and food patches frequently and spent considerable time in each (**Figure 2A**). We fit mixed-effects models to understand which factors affected the frequency and duration of patch visits (full model results reported in **Figure S1**).

**Figure 2.**
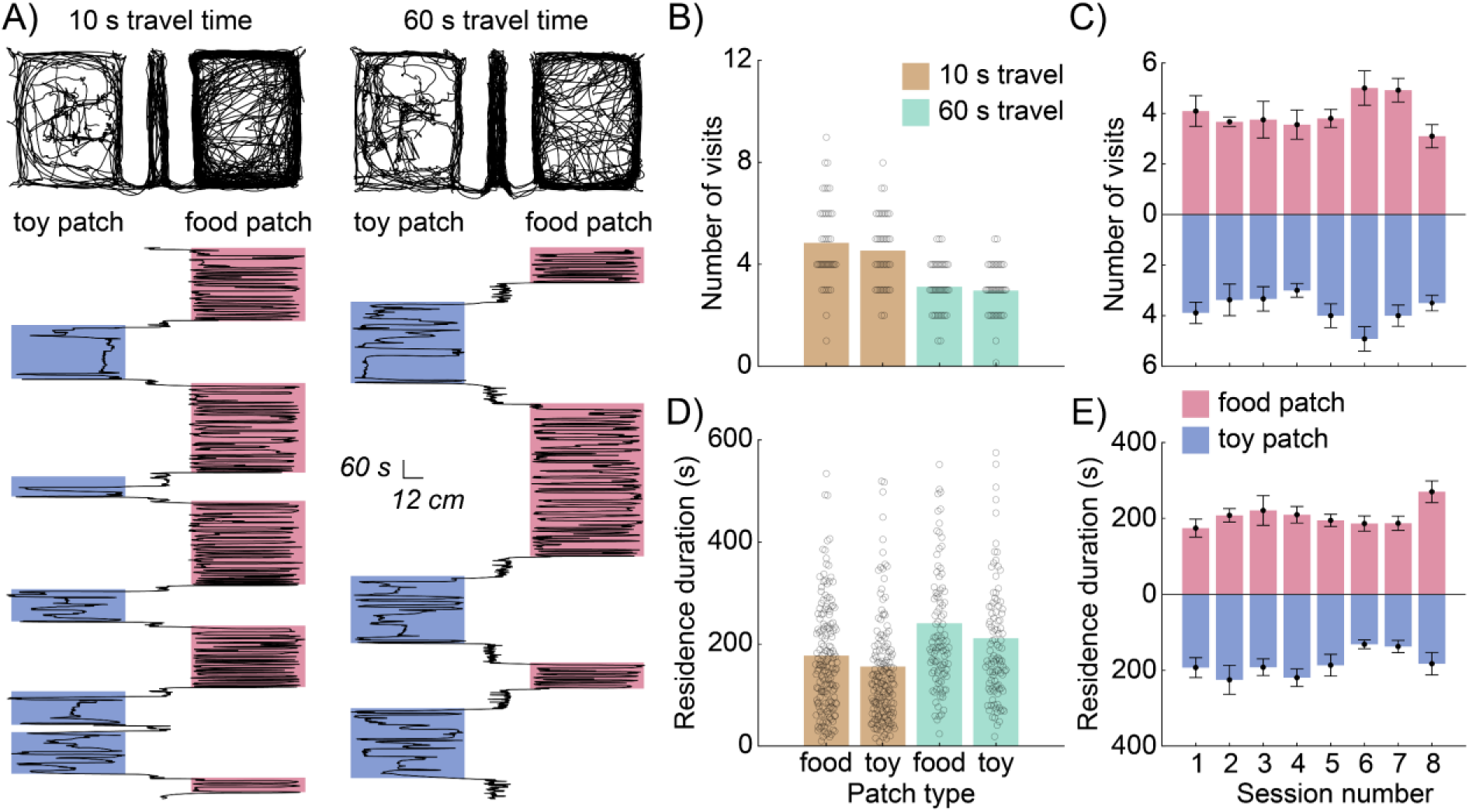
Behavioral results. (A) Position tracking data is shown for two behavioral sessions, one with a short (left) and one with a long (right) travel time. Plots at the top of the panel show top-down tracking data cumulated over the entirety of each session. In the bottom portion of the panel, the rat’s x-coordinate is plotted as a function of time within the session, with the start of the session at the bottom of the panel. Foraging bouts to toy and food patches are outlined with colored boxes. (B) Bars show the mean number of visits per session rats made to food and toy patches, separated by the long and short travel time conditions. Each data point indicates the number of visits rats made to each type of patch in an individual session (N = 160 patches in 80 sessions from 10 rats). (C) Bars depict the mean number of visits per session rats made to food and toy patches over the course of the eight-session testing sequence. Error bars depict the standard error of the mean. (D) Bars denote the average residence duration for patches of each type, separated by the long and short travel time conditions. Data points depict the residence time for a single patch within a single session (N = 160 patches in 80 sessions from 10 rats). (E) Bars display the mean residence time in food and toy patches for each of the eight sessions that comprised the testing sequence. Error bars depict the standard error of the mean.

Rats visited toy and food patches with equal frequency (**Figure 2B**; visit-number model; β_patch type_ = - 0.04; p = 0.74), but stayed ∼20% longer on visits to food patches (**Figure 2D**; visit-duration model; β_patch type_ = −0.21; p = 0.01). Travel time was a significant predictor of visit number (visit-number model; β_patch type_ = −0.22; p = 5.71×10^-7^) and visit duration (visit-duration model; β_travel time_ = 0.19; p = 8.57×10^-10^), with longer travel times resulting in increased patch residence duration, and thereby decreasing the number of visits to patches. This is consistent with models of optimal foraging, which suggest that visit duration should increase with the cost of seeking out a new patch (21, 33). The long travel time affected patch residence duration similarly for toy and food patches, increasing average visit duration in food patches by 36.0% compared to short travel time sessions, and increasing the average visit duration in toy patches by 35.3% relative to short travel time sessions. This suggests that the cost of switching foraging locations modulated decisions about food and non-food reward in a similar, opportunity-cost sensitive way.

We assessed whether preferences for food and toy patches were stable over time by including session number and its interaction with patch type as a predictor in both models. Residence duration was modulated by the interaction of session number and patch type (visit-duration model; β_patch type*session number_ = −0.14; p = 0.02), while the number of visits to patches was not (visit-number model; β_patch type*session number_ = 0.009; p = 0.91). To further investigate how session number and patch type interacted to affect residence time, we computed the correlation between residence time and session number separately for toy and food patches. Across sessions, rats subtly but significantly decreased the amount of time they spent in toy patches (R = −0.15; p = 0.02; **Figure 2E**). This may indicate that rats’ initial attraction to toy patches was partially driven by novelty, which decreased over time, or that rats learned to optimize the amount of time they spent in different patch types with experience, eventually settling into a preferred ratio of food and toy reward. The correlation between session number and residence duration in food patches was not significant (R = 0.07; p = 0.23; **Figure 2E**), nor was session number significantly correlated with the number of visits to food (R = 0.06; p = 0.60) or toy (R =0.13; p = 0.26) patches (**Figure 2C**). Thus, despite a subtle change in time allocation to toy patches across sessions, rats’ preferences were largely stationary.

### Movement patterns differed between food and toy patches

Rats could not initially know that food would never be available in toy patches, so it is possible that visits to toy patches reflected exploratory food seeking rather than utility derived from the toy. Because the play structure was consistently located in the center of toy patches, we used time spent in the center of the arenas as a proxy for rats’ engagement with the toy (**Figure 3A–B**). The amount of time rats spent in the center of patches was greater in toy patches compared to food patches (**Figure 3C**; β_patch type_ = 1.37; p = 6.82×10^-23^). In addition, rats traversed the center region of food patches quickly, but moved much more slowly through the center of toy patches (**Figure 3D**; β_patch type_ = - 1.78; p = 2.68×10^-59^), consistent with them pausing to engage with the play structure in toy patches, but moving swiftly through the center of food patches in pursuit of sucrose pellets.

**Figure 3.**
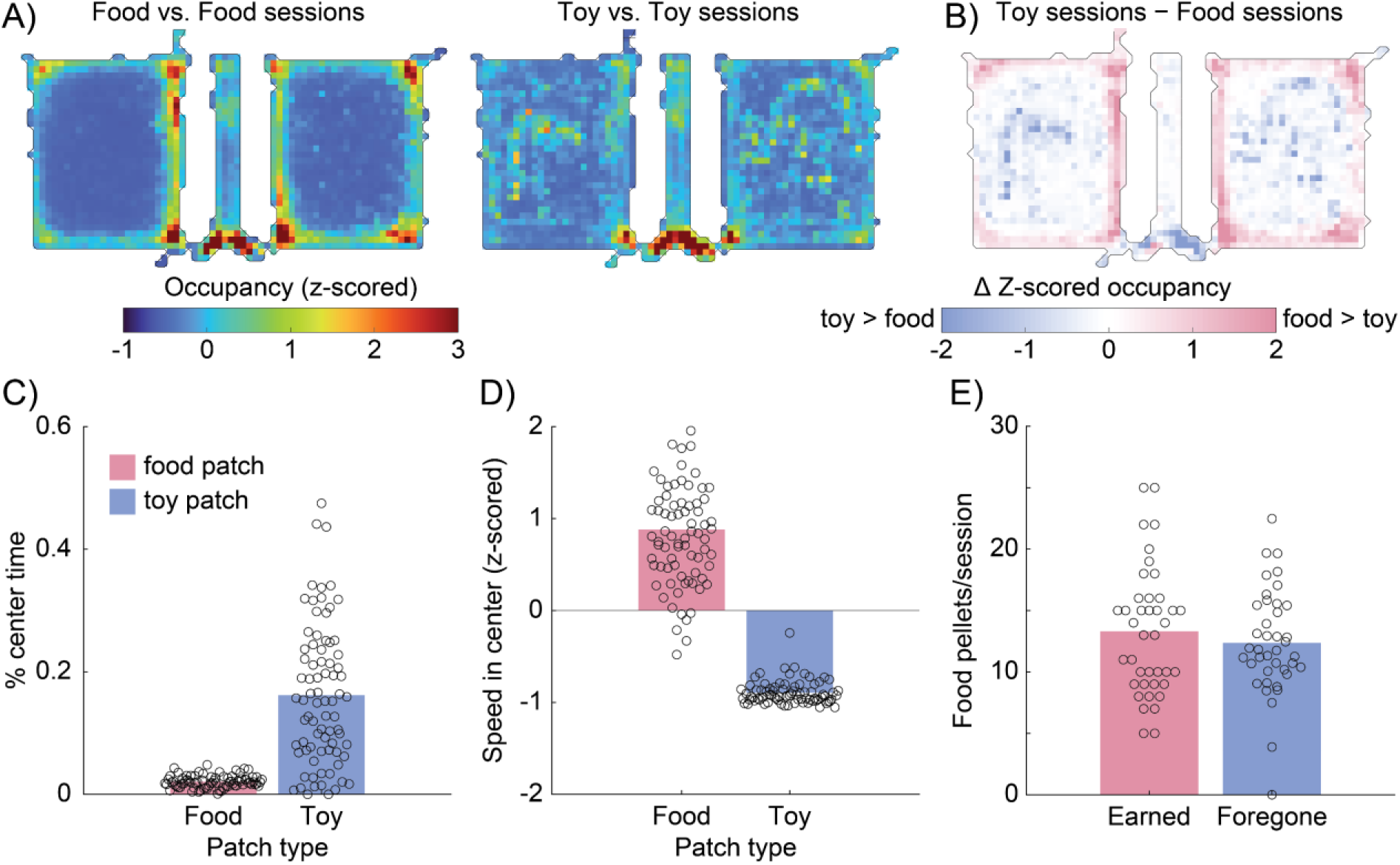
Rats engaged with the play structure, and visits to toy patches were costly. (A) The heat maps depict the average amount of time rats spent at each location in the apparatus in food vs. food session (left; N = 20 sessions from 10 rats) and toy vs. toy session (right; N = 20 session from 10 rats). Note the increase in time spent near the center of patches during toy vs. toy sessions, suggesting that rats actively engaged with the play structure. (B) The heat map plots the average difference in time spent at each location in the apparatus between toy vs. toy and food vs. food sessions. White colors indicate little or no difference in time spent between the two session types, while blue (red) colors denote more time spent in locations during toy vs. toy (food vs. food) sessions. Rats spent more time near the center of patches near the play structures in toy vs. toy sessions compared to food vs. food sessions. (C) We computed the fraction of time that rats spent in the center of patches during their visits to patches of either type (N = 160 patches in 80 sessions from 10 rats). Bars denote mean percentage of time spent in the center of food and toy patches, and data points show results from one patch in one session. Rats spend substantially more time in the center of patches when visiting toy patches. (D) We measured rats’ mean running speed in the center of patches and split the results by patch type (N = 160 patches in 80 sessions from 10 rats). Bars depict average center running speed for food and toy patches, while individual data points show average center running speed in one patch for one session. Rats moved quickly through the center of food patches, but moved much more slowly through the same portion of the apparatus in toy patches, consistent with rats pausing to engage with the toys. (E) For sessions in which rats choose between food and toy patches (N = 80 sessions from 10 rats) we compared the total amount of food they earned and the amount of food they could have earned on average by re-allocating their toy-patch visits to the food patch. The number of earned and foregone pellets was similar, indicating that visits to toy patches were costly in terms of lost opportunity to earn food.

### Visits to toy patches were costly

To quantify the cost rats paid by visiting toy patches, we focused our analyses on session types with one food and one toy patch (n = 40 sessions). In these toy-patch vs. food-patch sessions, the opportunity cost of spending time in the toy patch was clearly defined because visits to the toy patch could have instead been devoted to the food patch, where pellets were delivered according to the same depleting reward schedule (**Figure 1B**) on every visit. We computed the number of additional pellets rats would have earned on average had each toy-patch visit in a session instead been spent in the food patch (**Figure 3E**). The number of additional pellets rats could have earned in each session by only visiting the food patch was not significantly different than their actual earnings (β_earned/foregone_ = 0.14; p = 0.53). Thus, rats could have earned approximately twice as much food by avoiding the toy patch, indicating that visits to the toy patch came at a substantial cost in foregone food.

### Dynamics of food and toy patch visits

Finally, we tested whether rats had consistent preferences for the order in which they consumed food and non-food reinforcement. For instance, one might expect a food-restricted rat to visit food patches in the early part of sessions and allocate time to toy patches later, after its hunger was partially sated. In fact, we observed the opposite pattern (**Figure 4**). Examining how rats allocated time to the food and toy patches over the course of the session revealed that rats initially devoted more time to the toy patch, and later shifted their time allocation to favor the food patch (**Figure 4A**). We computed the latency for rats to accumulate 100 s, 300 s, and 500 s of time spent in each patch (**Figure 4B**) in toy-patch vs. food-patch sessions. Rats reached 100 s (β_patch type_ = −0.82; p = 1.27×10^-4^) and 300 s (β_patch type_ = −0.72; p = 0.001) of residence time more quickly for toy patches than food patches. Patch type did not affect latency to reach 500 s of residence time (β_patch type_ = −0.07; p = 0.76). This is consistent with time allocation to food patches eventually catching up with and overtaking time allocation to toy patches (**Figure 4A**).

**Figure 4.**
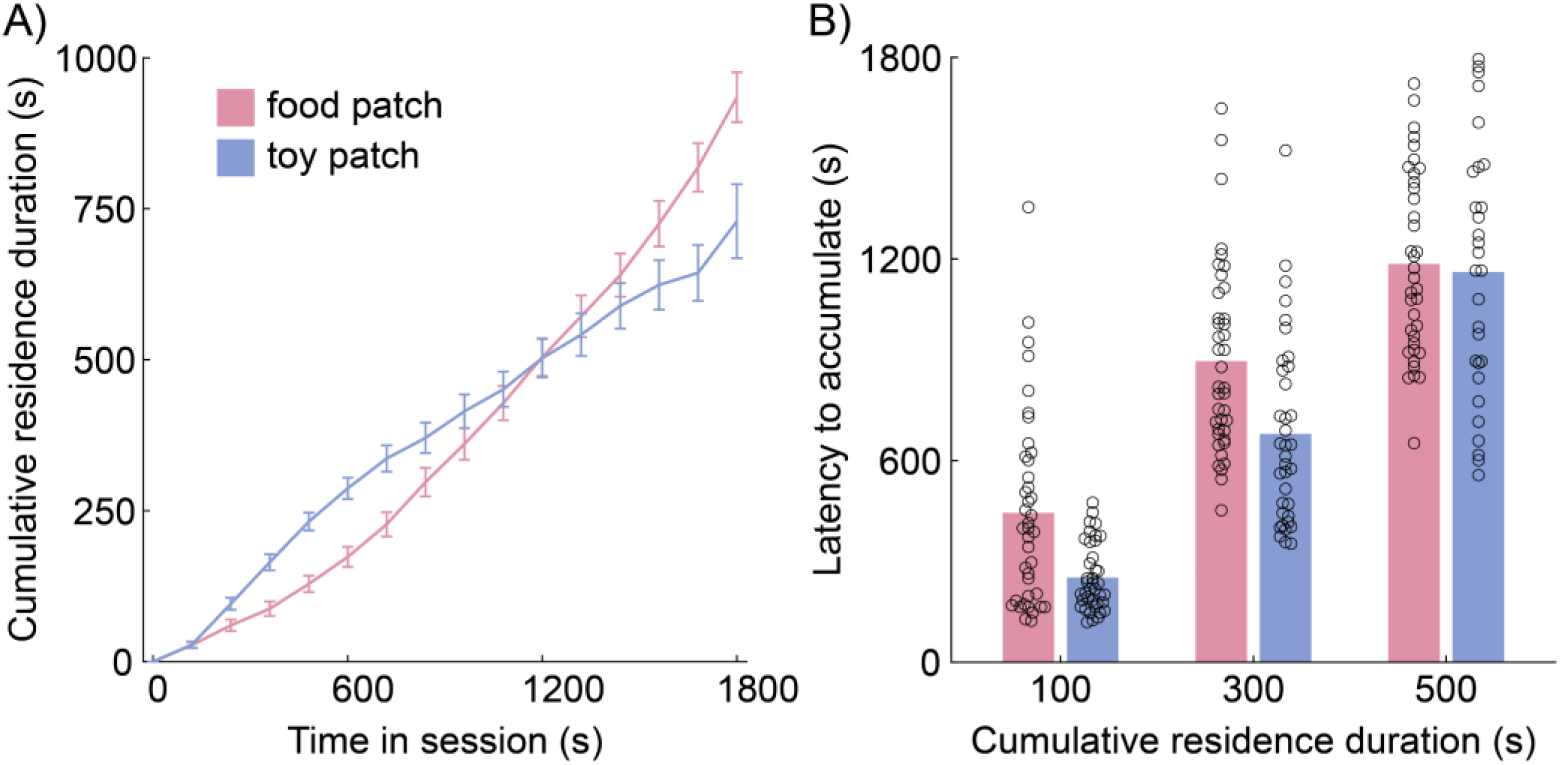
Temporal dynamics of food and toy patch visits. (A) For toy patch vs. food patch sessions, (N = 80 sessions from 10 rats) we tracked cumulative time spent in patches of each type across the 30-minute behavioral session. Early in sessions, rats devoted more time to toy patches, with the balance eventually shifting to favor food patches after around 2/3 of the session elapsed. (B) To quantify rats’ early preference for toy patches, we computed the amount of session time required for rats to accumulate 100, 300, and 500 s in patches of each type. The latency to reach 100 and 300 s was significantly lower for toy patches, but there was no difference in latency to reach 500 s in patches of either type, suggesting that rats initially preferred to devote time to the toy patch, but shifted to allocating longer-duration visits to the food patch in the final 1/3 of the session.

### Toy-patch visits persisted in the absence of food motivation

The results thus far suggest that rats derived utility from both toy and food patches and balanced their pursuit of both options during behavioral sessions. However, it is also possible that visits to toy patches were motivated by exploratory food seeking rather than interest in the play structure itself. Although rats never encountered food within toy patches, they may have continued investigating them to ensure that food did not unexpectedly become available. Such a strategy could be useful in environments where resource availability changes abruptly and unpredictably.

To address this possibility, a second cohort of rats (N = 6; 3 female) was tested on a sequence of sessions in which they chose between a toy patch and an empty patch, which contained neither toy nor food. For these sessions, the travel time was fixed at 10 s. Importantly, these rats always had *ad libitum* access to food in their home cage and never encountered any form of food within the testing apparatus. In this way, we sought to minimize both their motivation for seeking food during testing sessions and their expectation of finding food within the apparatus. We reasoned that if rats continued to devote time to the toy patch it was unlikely that they were doing so in search of food.

Under these conditions, rats showed an even stronger preference for the toy patch than in our initial experiments (**Figure 5A–B**). Rats made significantly more visits to the toy patch than to the empty patch (visit-number model; β_patch type_ = 0.59; p = 0.003), and visits to the toy patch were significantly longer, lasting on average twice the duration of visits to the empty patch (visit-duration model; β_patch type_ = 0.51; p = 0.001; full model results reported in **Figure S2**). This suggests that toy patches were attractive to rats in the absence of motivation for or expectation of food.

**Figure 5.**
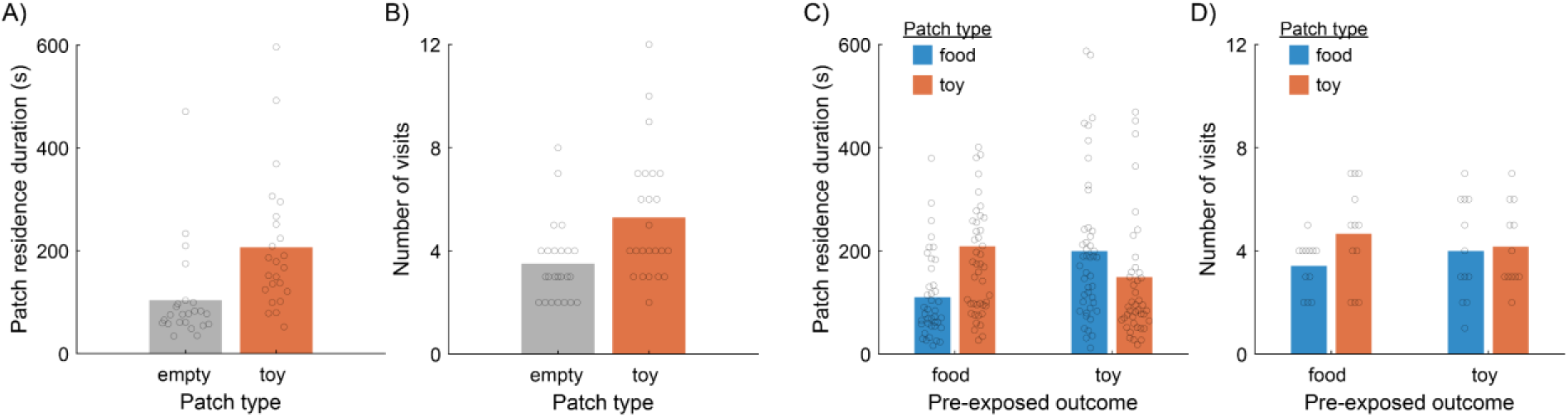
Motivational manipulations. (A) When choosing between a toy patch and an empty patch (N = 24 sessions from 6 rats), rats that were not food restricted made longer visits to the toy patch than the empty patch, suggesting that visits to the toy patch were rewarding and not simply driven by food seeking. (B) Rats made significantly more visits to the toy patch than the empty patch, confirming that both the frequency and duration of visits to the toy patch were enhanced relative to an empty patch. (C) Pre-exposure to food or toy prior to testing resulted in a decrease in time spent in the patch containing the pre-exposed outcome (N = 24 sessions from 6 rats). (D) The frequency of visits to food and toy patches was not affected by pre-exposure, suggesting that rats maintained a tendency to explore both patches following pre-exposure to one outcome, but allocated greater time to pursuing the outcome that was not made available prior to testing.

### Rats adjusted time allocation based on recent reward consumption patterns

Goal-directed agents that have recently consumed a particular reward—and thereby become partially sated on it— typically reduce their effort to obtain more of that outcome, indicating that their choices reflect current valuations rather than rigid habits. Thus, if rats aimed to balance their pursuit of food and non-food reward, inducing partial satiety for food should shift their time allocation to favor toy patches, and vice versa. To test this, the second cohort of rats was food-restricted for 7 days to induce hunger and then tested on a sequence of sessions in which they chose between a toy patch and a food patch, with the travel time fixed at 10 s. Prior to each testing session, rats were pre-exposed to either toy or food outside of the testing context to induce partial satiety for one of the rewards. Pre-exposure to food caused rats to decrease the duration of visits they made to the food patch, while pre-exposure to the toy caused rats to make shorter visits to the toy patch (**Figure 5C–D;** visit-duration model; β_pre-exposed outcome_= −0.41; p = 1.04×10^-4^). Pre-exposure did not affect the frequency of visits to either patch type (visit-number model; β_pre-exposed outcome_= −0.15; p = 0.27; full model results reported in **Figure S3**). Thus, when partially sated on one outcome, rats continued to sample both patches, but spent more time pursuing the reward that had not been provided before the testing session. This indicates that rats’ time allocation reflected their current valuation of the two reward types.

## Discussion

Using a free-choice foraging task, we found that hungry rats devoted a surprisingly large fraction of their limited time budget to seeking non-food reinforcement. Importantly, rats were sensitive to the opportunity cost of time in their pursuit of both food and non-food patches (20, 22, 33–35); this suggests that similar principles of rational decision making hold over a range goods, and may point to a common, integrated neural substrate that supports utility-maximizing decisions across qualitatively-distinct sources of reward. It has long been known that laboratory rats are motivated to explore unvisited spaces and investigate novel objects (36, 37); our data situate these findings in a quantitative, economic framework, enabling us to calculate the cost (in terms of lost food earnings) rats were willing to pay to engage with the play structure in toy patches. Rats in our experiment could have doubled their food earnings by not visiting toy patches, which suggests that the value they derived from interacting with toy objects was considerable.

There are at least two broad reasons that rats might have devoted time to toy patches. First, exploration of novel stimuli, spaces, or actions may have instrumental value by leading to the discovery of new sources of reward. In reinforcement learning (38) terminology, exploratory choices ensure that agents learn an accurate value function over their surroundings (39–43). Second, interacting with toy patches may have been an end in itself, just as many behaviors that humans engage in offer subjective satisfaction without easily measured instrumental benefit (44). The results of our satiety manipulation experiments suggest that both of these explanations factor into rats’ decisions. When rats were partially sated on one outcome before testing, they reduced the amount of time they spent during visits to the patch containing the pre-exposed outcome, but their frequency of visiting that patch was not affected. This suggests that time allocation was sensitive to recent consumption patterns, with rats decreasing their investment in a patch yielding an outcome they had recently received for free, while continuing to make brief visits to check whether the available outcomes had changed. These findings indicate that rats adaptively balanced exploration (continued sampling of both patches) and exploitation (devoting more time to the patch offering the scarcer, non-pre-exposed outcome) to maximize their utility.

Foraging models have increasingly been applied to decision neuroscience experiments (20, 45–47). Classic models of patch foraging maximize a single, intuitive currency: net rate of food intake (21, 22, 33, 48, 49). Such models would predict very short residence durations in toy patches since visits there never produced food. Although more complex foraging models have been developed to optimize over multiple currencies simultaneously (21), single-currency models remain in wide use and often make accurate behavioral predictions.

One well known systematic mismatch between model and data is the tendency of foragers to “overharvest” patches by staying longer than the optimal residence time (25, 26, 30, 50–53). Many explanations for overstaying have been proposed, and it seems likely that multiple factors contribute to this tendency (54–57). Our data raise the possibility that multi-objective optimization could play a role in overharvesting. Although rats in our experiment were hungry, they continued to value consumption of non-food rewards, and this is likely the case in experiments in which food reward is the sole programmed reinforcer. Overstaying in such settings might be driven by foragers deriving satisfaction from whatever alternative behaviors are available to them within a patch, even if such opportunities are minimized by the experimental design. A challenge for future research is to better characterize the currency that foragers aim to maximize, which may be complex even in relatively constrained behavioral contexts.

Future work could further test whether rats’ foraging decisions comport with additional principles of rationality. For instance, increasing or decreasing the rate of reward in food patches offers another means of varying the opportunity cost of spending time in the toy patch; a higher rate of reward might shift time allocations to further favor the food patch, while a decrease in food delivery rate would be expected to drive additional time allocation to toy patches. Parametric variation of non-food reinforcer options could be used to further fractionate rats’ preferences and provide more detailed insights into how rats derive utility from engaging with different types of objects. More generally, the free-choice foraging behavioral framework used here could be extended to investigate a wider range of both nutritive (58, 59) and non-nutritive (60–63) rewards.

## Methods

### Subjects

Long-Evans rats (n = 16; 8 female) were purchased from Charles River at approximately 300 g, and were maintained on a standard 12-hour light cycle. Rats were housed singly and when food restricted were maintained at 85% of their free-feeding weight. Rats earned food reward in the behavioral task, but our experiments did not follow a “closed economy” design, so supplemental rodent chow was provided daily to ensure they maintained a stable weight. Rats were handled 10– 15 minutes per day for a week before behavioral testing commenced. During the week of handling, rats were habituated to the behavioral apparatus by allowing them to explore it for 10 minutes per day with all doors open. Before being tested in the experiments described here the first cohort of 10 rats completed 24 sessions of a similar foraging task in the same apparatus; these sessions involved food-patch vs food-patch comparisons exclusively. Rats never experienced the toy option prior to the experimental sessions reported here. All procedures were approved by the Chancelor’s Animal Research Committee at the University of California, Los Angeles.

### Behavioral task

Rats were tested once per day in sessions lasting 30 minutes. In each session, rats foraged in an automated 3-chamber apparatus with 80 cm walls. Rats were placed in the travel corridor (18×71 cm) at the beginning of each session. After the travel delay elapsed, the doors blocking access to patches (61×71 cm) opened simultaneously, allowing rats to choose which location to visit. Upon entering one patch, the door to the unchosen patch closed. Sucrose pellets (45 mg; Bioserv) were delivered from dispensers (Med Associates) mounted above each patch. Dispensers were positioned such that the pellets scattered unpredictably and approximately uniformly across each arena. The travel time delay was either 10 s or 60 s, and remained fixed for the duration of the session.

The initial reward rate in food patches was 1 pellet every 26 seconds. The reward rate decreased exponentially (τ = 90 s) while the rat remained in the patch, and reset to its maximal value each time the rat re-entered a food patch following a travel time delay. The timing of pellet deliveries was stochastic, approximating an inhomogeneous Poisson process, and a unique set of pellet delivery times was programmed for each food patch visit using a sequential procedure. The latency to the first pellet delivery was drawn from an exponential distribution with λ set to 26 s (i.e., an inter-pellet interval consistent with the maximum rate of reward). Subsequent inter-pellet intervals were similarly drawn from exponential distributions with λ set to the inter-pellet interval corresponding to the patch’s current reward rate. The mean inter-pellet interval after *t* seconds in the patch was computed as:

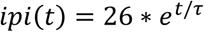

In toy patches, rats encountered a commercial “hamster playground” (Wontee) containing beams for climbing and fixed strings suspending metal rings and small pieces of wood. The footprint of the playground was 30 cm × 20 cm, and it was 18 cm tall at its highest point. The playground was placed in the center of the patch and was too heavy for rats to move. Rats were free to interact with these objects as they chose to. Rats’ location was tracked during behavioral sessions using Bonsai (64). Reward deliveries and door positions were controlled by custom Matlab (version 2024a, Mathworks) code. The play structures and the entire apparatus were cleaned thoroughly with 70% ethanol after each behavioral session.

The first cohort of 10 rats were tested on an eight-session sequence that exhaustively assigned patch type (toy or food) to both locations within the apparatus (left or right) at two levels of travel time (10 s or 60 s). The patch-type assignments were therefore: toy-left/toy-right, food-left/food-right, food-left/toy-right, and food-right/toy-left. These combinations were repeated once with the long travel time and once with the short travel time. Rats experienced session types in random order.

The second cohort of 6 rats were tested in a two part experiment. First, prior to experiencing any food restriction, rats were tested on a sequence of sessions comparing a food patch with the same reward statistics as above, and an empty patch that did not contain a toy structure or deliver sucrose pellets. Patch type assignments were: toy-left/empty-right, and empty-left/toy-right. Each combination was repeated twice per rat, with travel time fixed at 10 s, resulting in a total of 4 sessions per rat. Rats experienced session types in random order. After completing these sessions, rats were placed on food restriction for 7 days to reduce their body weight to 85% of their free-feeding weight, and then proceeded to the satiety manipulation experiment. Here, rats were tested on a sequence of comparisons between a toy patch and a food patch, but before each session they were pre-exposed to either the toy structure or 15 g of sucrose pellets for 30 minutes in an open-field that was distinct from the testing apparatus. Patch type assignments were: toy-left/food-right, and food-left/toy-right, and these were fully crossed with toy and food pre-exposure, resulting in a total of 4 sessions per rat. Travel time was again fixed at 10 s for all sessions. Rats experienced session types in random order.

### Data analysis and statistics

All data analyses were completed using Matlab 2024a. We used linear or generalized linear mixed-effects models to quantify most aspects of behavior on the task (65). Two main models tested which factors affected the number of visits rats made to each patch within a session (generalized linear mixed effects model with Poisson link function to accommodate count data; Matlab function *fitglme*), and the amount of time rats spent in a patch on each visit (linear mixed effects model; Matlab function *fitlme*). We included the same set of predictors for both models (**Figure S1**). Continuous predictors (travel time and session number) were z-scored to standardize coefficient estimates and facilitate comparison of effect sizes. Categorical predictors were sex (female coded as one), patch type (toy patch coded as one), and the location of the patch within the apparatus (side; left coded as one). Rat identity was included as a random effect. The residence duration of each visit was defined as the time between when rats first crossed from the travel corridor into one of the two patches, to when rats re-entered the travel corridor far enough to trigger the doors to close and initiate the travel time delay. To remove skew in residence time data and improve model fit, residence times were log transformed. If the session ended while a rat was still foraging in a patch the final visit was excluded from analysis of residence durations, due to uncertainty over how much longer the rat would have remained in the patch. To assess the effect of reward pre-exposure in the satiety experiment, we created a categorical predictor that tracked whether the identity of the pre-exposed outcome was the same as the reward type available in each patch. For mixed effects models quantifying other aspects of behavior, the predictors included are described in the main text; in all of these models, rat identity was included as a random factor.

To visualize the distribution of behavior across the patches, we divided the apparatus into a 100×50 grid of bins, and computed the amount of time rats spent in each bin during each session. Occupancy at each location was z-scored within session and then averaged across sessions, splitting by patch type to produce the heat maps in **Figure 3A**. The occupancy difference heat map in **Figure 3B** was produced by subtracting pixel-by-pixel the average heatmap of toy vs. toy sessions from the average heat map of food vs. food sessions. For analyses of time spent and running speed in the center of patches, the rat was counted as occupying the center region of a patch when their distance to the center of the patch was 17 cm or less. Choosing different values for this threshold did not affect the qualitative or statistical pattern of results.

We defined the opportunity cost of toy-patch visits as the number of additional food pellets rats could have earned had every toy-patch visit been allocated to the food patch instead. Reward rate in food patches declined over time in the same way on each visit, but stochasticity in the timing of pellet deliveries meant that two visits of the same duration did not necessarily result in the same number of pellets being dispensed. Accordingly, to find the opportunity cost of toy-patch visits, we simulated pellet delivery times for 1000 patch visits as described above, and computed the average number of pellets that would have been delivered over the duration of each toy patch visit.

## Acknowledgements

This work was supported by Whitehall foundation research grant #2021-12-045 (A.M.W.) and 1R01MH137276 (A.M.W.). R.L. and M. D. were supported by the UCLA-HBCU Neuroscience Pathways program. The authors thank members of the Wikenheiser lab for helpful discussions about this work.

## Data availability

Data and code are available at https://doi.org/10.5281/zenodo.14481737.

---

**Figure S1.**
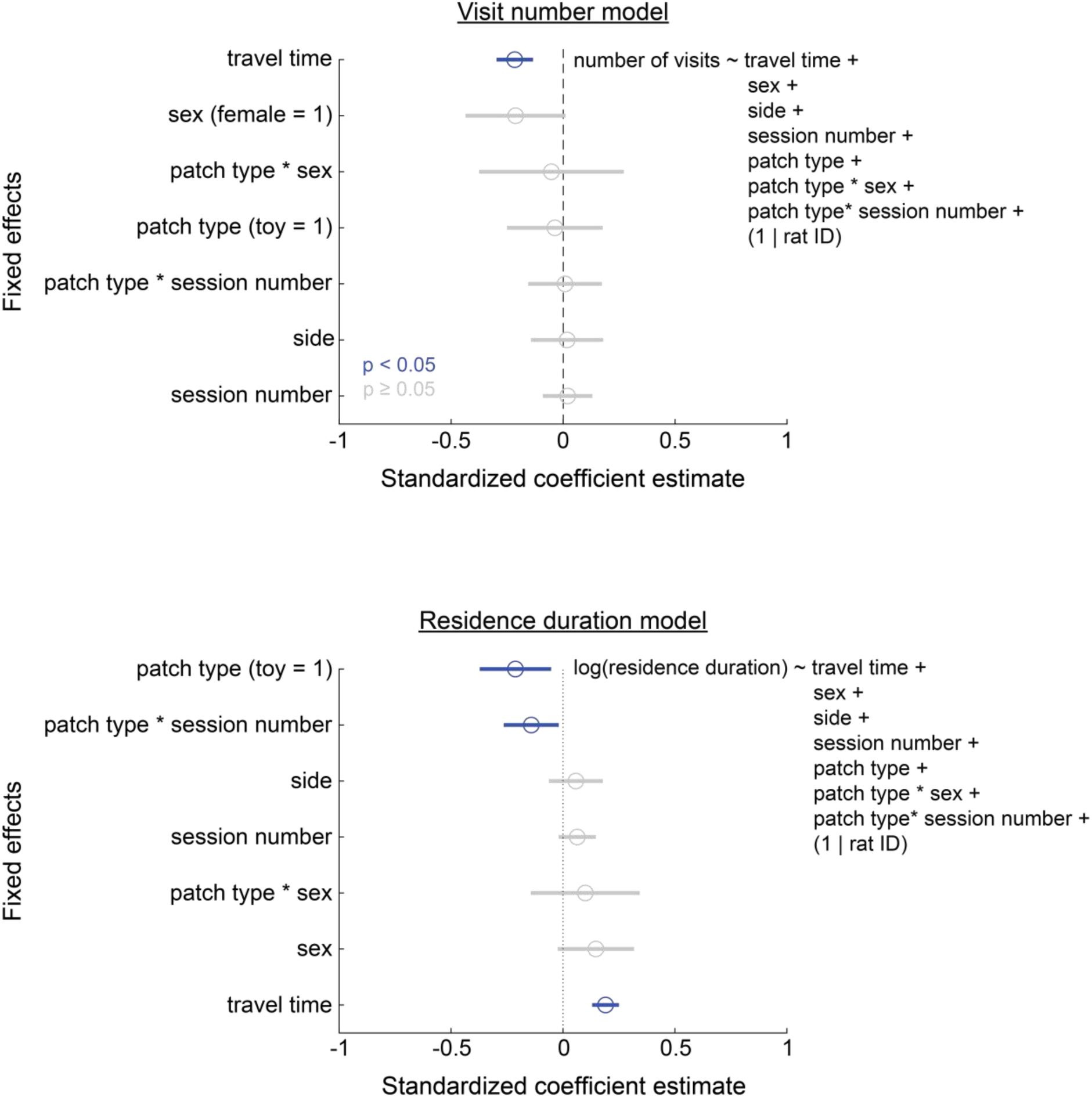
Regression model results: main data set. We used mixed-effects models to quantify which factors influenced the number and duration of visits rats made to patches. Coefficient estimates for each model are sorted by their magnitude, with significant predictors plotted in blue. Error bars indicated the 95% confidence interval around each coefficient estimate.

**Figure S2.**
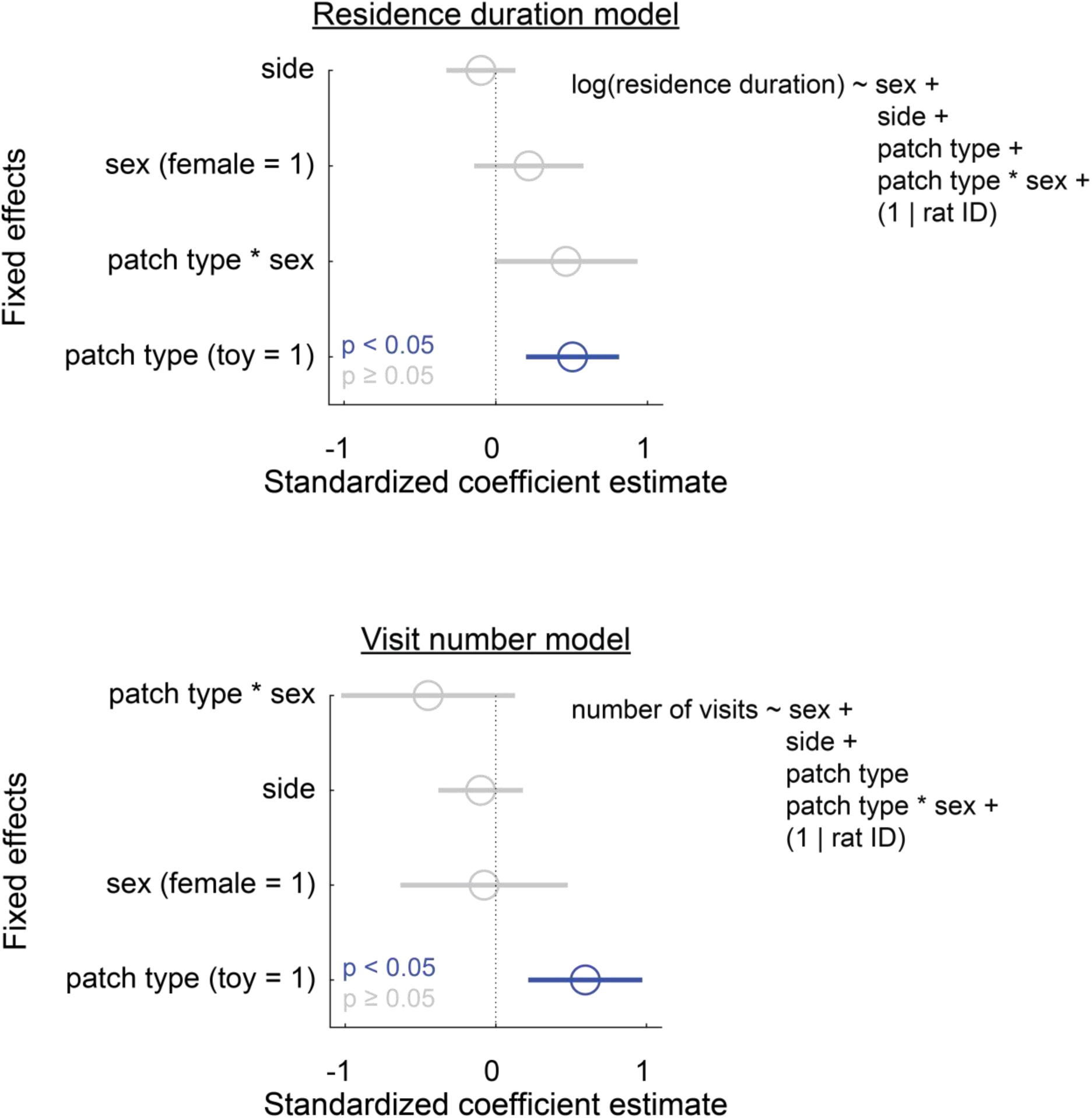
Regression model results: toy patch vs. empty patch sessions. Mixed-effects model quantification of factors that influenced the duration and frequency of visits rats made to toy and empty patches. Coefficient estimates for each model are sorted by their magnitude, with significant predictors plotted in blue. Error bars indicated the 95% confidence interval around each coefficient estimate.

**Figure S3.**
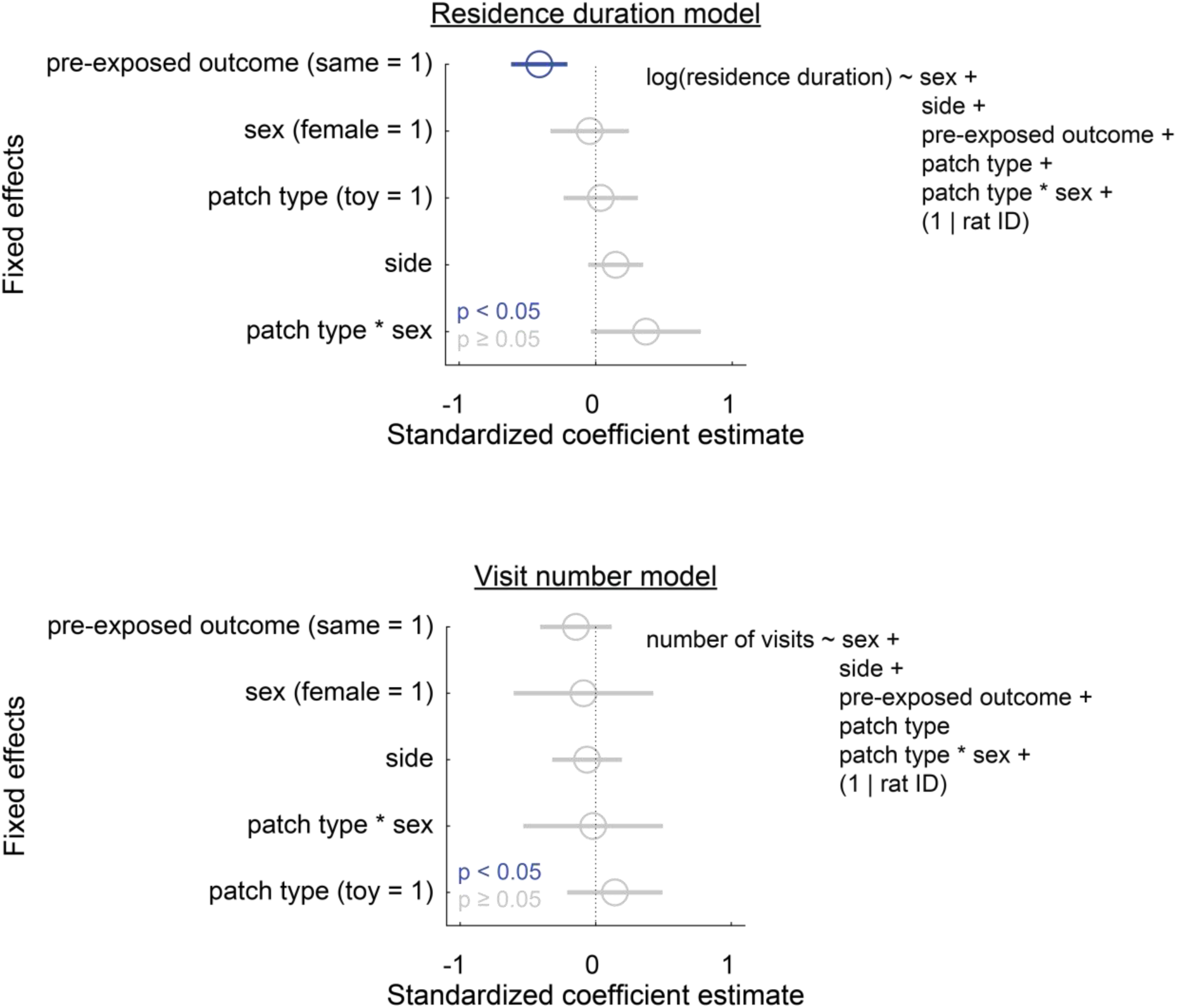
Regression model results: satiety manipulation sessions. Mixed-effects model quantification of factors that influenced the duration and frequency of visits rats made to toy and food patches following pre-exposure to one of these outcomes before the testing session. Coefficient estimates for each model are sorted by their magnitude, with significant predictors plotted in blue. Error bars indicated the 95% confidence interval around each coefficient estimate. The predictor “pre-exposed outcome” is one for patches that provided the same outcome as the rat received during pre-exposure and zero otherwise.

## Notes

### Competing Interest Statement

The authors have declared no competing interest.

### Summary of Updates

Manuscript revised to include new data.

